# Proteochemometric modeling strengthens the role of Q299 for GABA transporter subtype selectivity

**DOI:** 10.1101/2024.08.13.607728

**Authors:** Stefanie Kickinger, Anna Seiler, Daniela Digles, Gerhard F. Ecker

## Abstract

Proteochemometric modeling (PCM) combines ligand information as well as target information in order to predict an output variable of interest (e.g. activity of a compound). The big advantage of PCM compared to conventional Quantitative Structure-Activity Relationship (QSAR) modeling is, that by creating a single model one can not only predict the affinity of a diverse set of compounds to a diverse set of targets, but also extrapolate the specific ligand-protein interactions that might be relevant for activity. In this study, we compiled a dataset of 323 compounds and their bioactivity data regarding the inhibition of the four GABA-transporter (GAT1/BGT1/GAT2/GAT3) subtypes, which are potential new drug targets for treating epilepsy. Proteochemometric modeling using partial least squares and random forest provided models which performed equally well than conventional QSAR models for each individual transporter. However, by analyzing the importance of the protein descriptors used in the PCM models, we identified the amino acid Leu300/Q299/L294/L314/ in GAT1/BGT1/GAT2/GAT3 to be relevant for binding and subtype selectivity.

## Introduction

γ-Aminobutyric acid (GABA) is the principal inhibitory neurotransmitter in the central nervous system. Its inhibitory effect is mainly induced by ligand-gated GABA_A_-receptors, which are located synaptically initiating phasic inhibition, or extra synaptically causing persistent tonic inhibition^1^. Triggered by depolarization of the presynaptic axon terminals, GABA is released into the synaptic cleft and binds to postsynaptic receptors, thereby changing the electrical excitability of the postsynaptic neuron^2^. GABA transporters (GATs) terminate the GABAergic signaling by transporting GABA together with chloride and sodium from the synaptic cleft into presynaptic neurons and surrounding glial cells. Irregularities in this GABAergic neuronal system are known to be involved in various neurological diseases such as epilepsy, stroke, or Alzheimer’s disease^3^. Inhibiting GATs is proven to increase the level of GABA in the synaptic cleft and to be effective against seizures in epileptic disorders^4^.

In human and rat, four different GATs have been identified, which all belong to the solute carrier 6 (SLC6) family, and are labeled GAT-1 (SLC6A1), BGT-1 (SLC6A12), GAT-2 (SLC6A13), and GAT-3 (SLC6A11). In mouse, the same transporters correspond to GAT1-4, respectively^5^. Due to the somewhat confusing nomenclature in different species we will refer in this manuscript to the nomenclature according to the International Union of Basic and Clinical Pharmacology (IUPHAR) GAT1 (SLC6A1), BGT1 (SLC6A12), GAT2 (SLC6A13) and GAT3 (SLC6A11)^6^. Different species will be indicated by the lower case letters h (human), m (mouse) and r (rat).

Due to the different functions and different brain expression patterns, considerable effort has been spent on the development of GAT subtype-selective inhibitors. So far, highly selective and potent inhibition has been only accomplished for GAT1^7^. This might be due to the fact that GAT1 shares only about 47–49% sequence identity with the other GATs, whereas the non-GAT1 subtypes share about 63–70% sequence identity^5^. Recently, a cryo electron microscopy structure of human GAT1 in complex with the antiepileptic drug tiagabine was published^8^. Furthermore, several structural homologs such as the bacterial leucine transporter^9^ as well as other SLC6 family members e.g. the human serotonin transporter^10^ have been resolved and were successfully used as templates for homology models^11^. Docking studies of known ligands utilizing these homology models revealed the first insights into what is driving GAT subtype selectivity^12,5^. Nevertheless, there is still a lack of computational models which would allow to predict the interaction profile of a set of compounds with all four GABA transporter subtypes. Proteochemometric (PCM) modeling is a Quantitative-Structure-Activity-Relationship (QSAR) approach that utilizes ligand as well as protein information and thus allows to extrapolate relevant protein-ligand-interactions for binding^13^. This technique was already successfully applied to predict novel ligands for the SLC transporters SLC5A1 (Sodium-Glucose transporter 1, SGT1) and SLC5A2 (SGT2)^14^. In this study, we compiled and curated a dataset of 323 ligands inhibiting mGATs from the publically available data sources Chembl25^15^ and Pubchem^16^ and developed a set of proteochemometric models. Descriptor importance analysis revealed that the protein descriptors for the corresponding residues L300/Q299/L294/L314 and Y60/E52/E48/E66 (GAT1/BGT1/GAT2/GAT3, respectively) were ranked highest among the protein descriptors. Thus, we conclude that proteochemometric modeling is a promising computational technique capable of predicting protein-ligand interaction profiles of GABA-transporter.

## Methods

### Data set

In order to generate a comprehensive dataset without the need for manual extraction we generated a set of KNIME Analytics Platform 3.7^17^ workflows which extracted data from ChEMBL25^15^ and PubChem^16^, standardized the chemical structures and calculated a set of descriptors. In a first step, we retrieved all bioactivity data available for the four GATs in mouse, rat, and human (Uniprot IDs^18^: P30531, P31648, P23978, P48065, P31651, P48056, Q9NSD5, P31649, P31646, P48066, P31650, P31647).

All retrieved compounds were standardized according to the same protocol as already described by Girardi *et al*.^19^. Briefly, all compounds were neutralized and the representation of aromatic rings, double bonds, hydrogens, tautomers and mesomers were standardized. Stereochemistry information was removed to allow the identification of compounds that would be handled as duplicates by the 2D descriptors used. Subsequently, bioactivity values were transformed to their pIC_50_. Finally, the compounds were aggregated according to their InChIkeys to remove duplicate entries. To work with a homogeneous set of data points, we used only mouse GATs with IC50 as endpoint for our final dataset, as this provided the highest number of available compounds measured in the same organism (ChEMBL: 379 compounds tested at least in one GAT resulting in 1088 bioactivity entries, PubChem: 337 compounds, 923 bioactivity entries, see Figure 1). We further filtered the ChEMBL data to only contain entries annotated with “=” to ensure high data quality for subsequent regression modeling (removal of 50 compounds and 163 bioactivity entries). As PubChem does not provide a similar annotation, we removed bioactivity entries that showed a pIC_50_ of 3 (1000 μM) and 2.301 (5000 μM) since manual curation revealed that these entries were annotated with “>” in the original publications (removal of 58 compounds, 191 bioactivity entries). In case of multiple annotated bioactivities for the same compound-transporter pair, the arithmetic mean and the standard error of the mean (SEM) were calculated. Compound-transporter pairs that comprised a SEM bigger than 0.2 were considered as ambiguous and thus deleted from the dataset (removal of 32 compounds/40 bioactivity entries from ChEMBL data, 33 compounds/40 bioactivity entries from PubChem). Ultimately, we merged the ChEMBL (309 compounds, 699 bioactivity entries) and PubChem data (292 compounds, 692 bioactivity entries) data and retrieved a dataset containing 317 compounds tested at least in one GAT and 744 bioactivity entries with a fairly homogeneous activity distribution (Figure 1).

**Figure 1.**
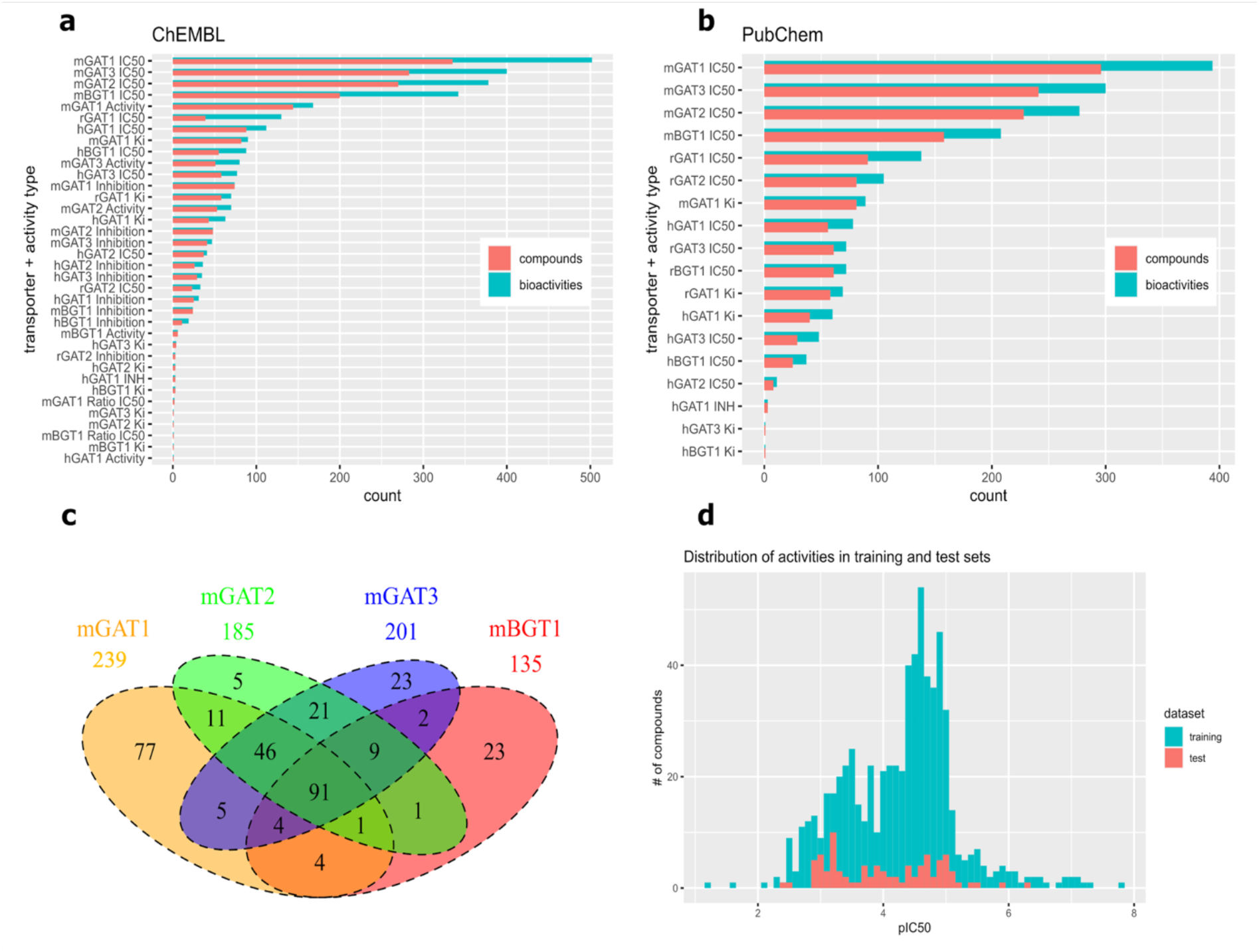
**a**. Distribution of available bioactivities/compounds per transporter, species and activity endpoint in ChEMBL before data curation. **b**. Distribution of available bioactivities/compounds per transporter, species and activity endpoint in PubChem before data curation. **c**. Venn diagram of the final curated GAT dataset including test and training set (in total 323 unique compounds and 760 bioactivities). **d**. Distributions of activities in the training and in the test set over all GATs.

The data was split into a training and test set by randomly selecting 16 compounds that were tested in all four GATs for the test set. Additionally, we added six compounds tested in all GATs from an inhouse dataset from Vogensen *et al*.^20^ to the test set. The test set ultimately comprised 22 compounds and 88 bioactivities. The training set comprised 301 compounds tested at least in one GAT (in total 672 bioactivities entries). Data Visualization was performed with KNIME 3.7 (“GroupBy Bar Chart node”) and in Rstudio 1.1.456 with R 3.6.3 and the packages ggplot2 3.3.0, corrplot 0.84 and VenDiagram 1.6.20. A Murcko Scaffold analysis was performed with the KNIME node “RDKit Find Murcko Scaffolds” (121 scaffolds were detected in total, 117 in the training set, 16 scaffolds in the test set, 12 scaffolds were present in both the training and the test set).

### Calculation of Descriptors

To describe the compounds’ properties, 25 physicochemical 2D-descriptors were calculated using the RDkit node in KNIME 3.7. To describe the protein properties, Z3-scales^21^ descriptors were calculated with the camb package^22^ in R 3.6.3 for the orthosteric pocket. The orthosteric pocket was defined as all corresponding GAT residues (15 in total) that are within a distance of 4.5 Å of the co-crystallized ligand leucine in the homologous LeuT crystal structure PDB ID 2A65 according to an alignment by Kickinger *et al*.^5^. This pocket is fully conserved in the different species of the respective transporter. As outlined in the alignment in Figure 3, only amino acids at the position 1, 2, 11, 12, 13, and 15 showed variance, mostly due to differences in GAT1. All protein descriptors with zero variance were removed from the dataset resulting in 18 protein descriptors (six different amino acid positions described by three z-scale descriptors each). To analyze the chemical space covered by training and test set we performed principal component analysis in R with the packages factoextra 1.0.7 and FactoMineR 2.3 (see Figure 2).

**Figure 2.**
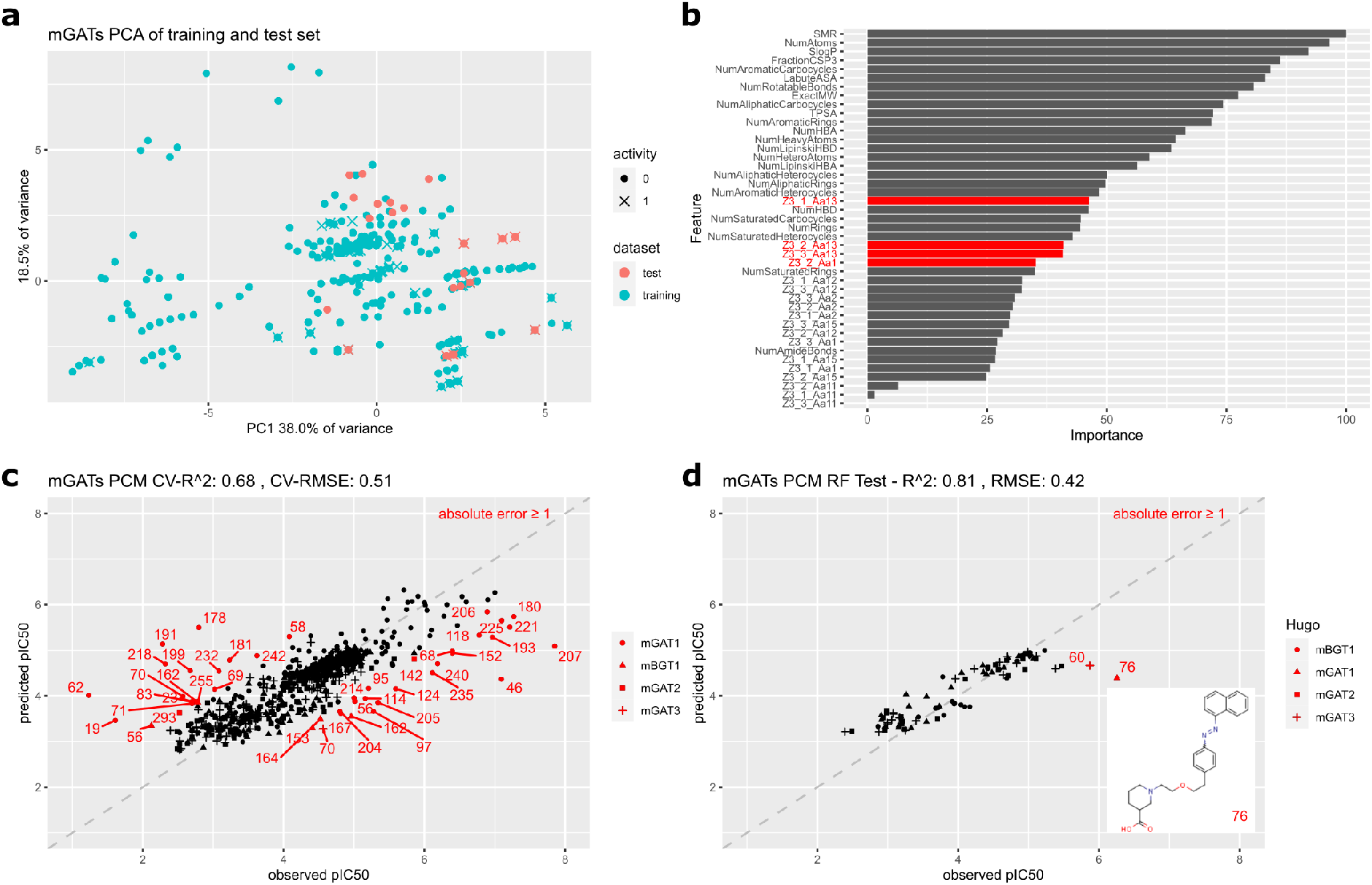
**a**. Principal component analysis (PCA) of the training and test set of all GATs based on 25 physicochemical RDKit descriptors; **b**. Importance ranking of descriptors for RF PCM model scaled to 100; **c**. 10-fold cross-validation performance of RF PCM model; **d**. Test set performance of RF PCM model and chemical structure of compound **76**.

**Figure 3:**
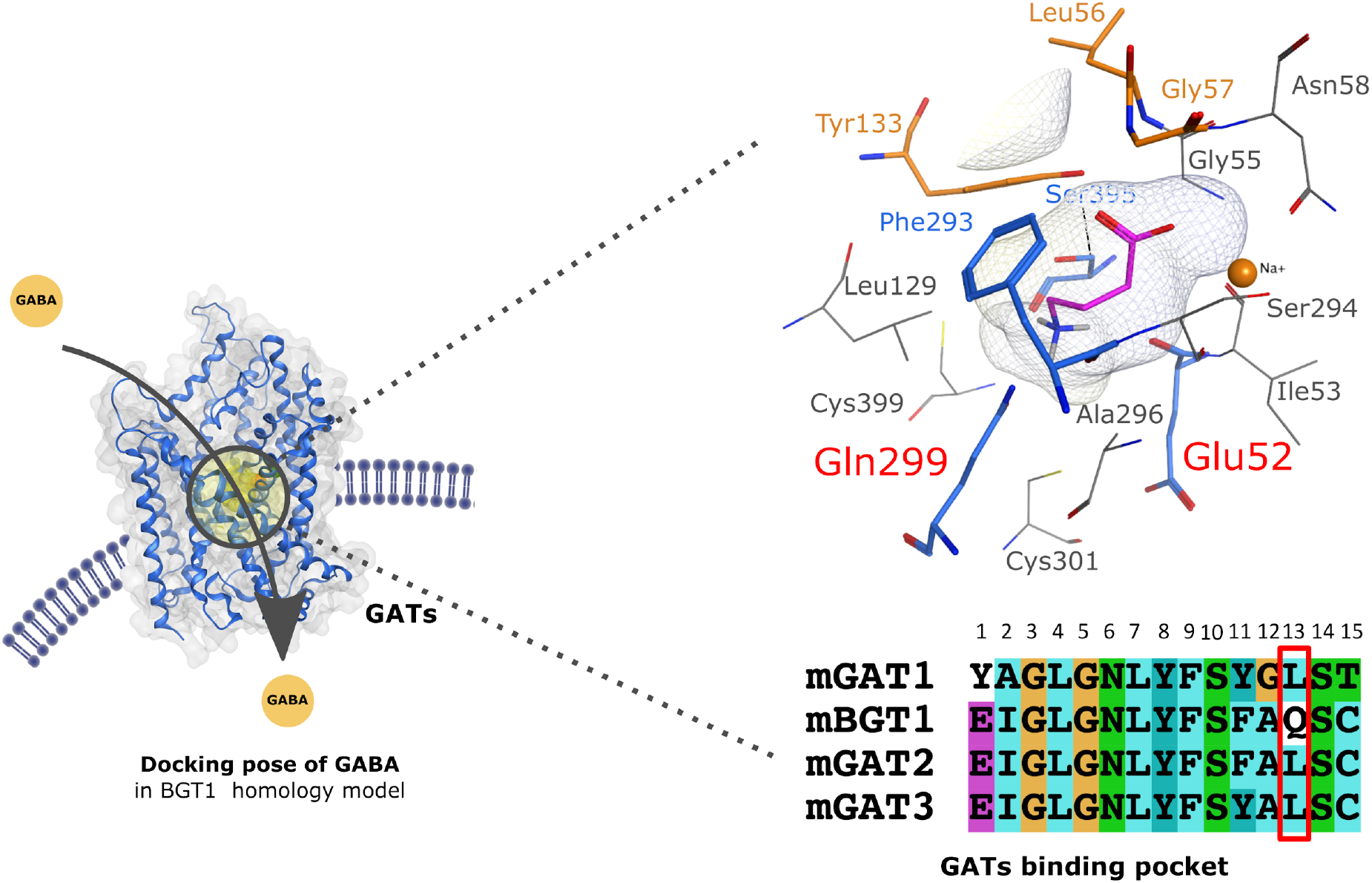
Docking pose of GABA in a homology model of hBGT1^5^. Residues in blue interact with the baroxal moiety of GABA, residues colored in blue interact with the amino moiety of GABA. Residues labeled in red have been identified by the RF PCM model as relevant.

### Model building

For model training, we used the caret 6.0-86 package in R 3.6.3, and trained Partial Least Square (PLS) as well as random forest (RF) models. For the PLS models we applied default options and automatically selected the number of components with the lowest Root Mean Square Error (RMSE). For the RF models we also used default parameters such as five features and 500 trees. The performance of the models was assessed by 10-fold cross-validation and by the prediction of a test set (calculations of R^2^ and RMSE). Compounds were considered as poorly predicted if the difference between the predicted pIC_50_ and the actual pIC_50_ was bigger than 1 (tenfold difference). Descriptor importance analysis was performed with the function varImp() in the caret package and was used to identify the most relevant ligand-based features as well as the amino acid position contributing the most to the models’ predictivity.

### Applicability Domain

The applicability domain of all generated models was calculated with the “Domain Similarity” node from the Enalos nodes for KNIME. The Euclidean distance of a test compound to its nearest training compound neighbour was calculated and compared with a predefined applicability domain threshold (default 0.5).

### Results and Discussion

Proteochemometric modeling (PCM) combines ligand information as well as target information in order to predict an output variable of interest (e.g. activity of a compound). The big advantage of PCM compared to conventional QSAR modeling is, that by creating only a single model one can not only predict the affinity of a diverse set of compounds to a diverse set of targets, but also extrapolate the specific ligand-protein interactions that might be relevant for activity. Since PCM has already been successfully applied to other SLC transporters^23,14,24^, we wanted to investigate whether PCM is a potential new method to predict the activity and selectivity of compounds inhibiting neurotransmitter transporters such as the four GABA transporters. Specifically, we were interested in two main questions: (i) how does a PCM model perform compared to a set of individual models calculated for each transporter in predicting the activity profile of new compounds, and (ii) can the PCM model identify amino acids relevant for activity and selectivity.

One of the most crucial tasks for creating predictive and robust machine learning models is ensuring sufficient data size and quality. The availability of large public data sources such as ChEMBL and PubChem, together with workflow tools e.g. KNIME, allow the creation of large datasets without the need of manual extraction from individual publications. With the help of KNIME workflows we retrieved all available bioactivity data from ChEMBL and PubChem for mouse, human and rat GATs. Analysis of the retrieved data showed that most data was available for mouse GATs annotated with the activity endpoint IC_50_ (see Figure 1a,b). Because of the availability of a high number of compounds tested in one species under similar assay conditions, we decided to build regression models for the available mGAT data. To ensure high data quality, which is essential for building robust regression models, we applied a rigorous data curation protocol (see Method section). The finally retrieved dataset comprised 323 unique compounds that were tested at least in one mGAT (see Figure 1c) resulting in 760 bioactivity entries, which served as basis for our proteochemometric modeling approach.

For calculating the PCM models, the dataset was split into a training set with 301 compounds (672 bioactivities) and a test set with 22 compounds tested in all GATs (88 bioactivities). For the individual models we used the available data for each transporter for the training set (mGAT1 217, mBGT1 113, mGAT2 163, mGAT3 179 compounds) as well as the same test set as for the PCM models. To analyse the diversity of the training and test set we performed PCA and plotted the first two dimensions (Figure 2a). As we restricted the generation of the test set to include only compounds that were tested in all mGATs in order to use the same test set for validating the PCM and the individual models, the test set does not cover the whole chemical space of the training set as shown in the PCA plot in Figure 2a. However, an applicability domain for the test set was calculated based on Euclidean distance which showed that the test compounds are within the domain of the training sets. The PCM models and the individual models were calculated using both PLS and RF. In all cases, RF performed considerably better than PLS with respect to 10-fold cross validation of the training set and the prediction of the test set. Consequently, only the RF models will be discussed further. As shown in Table 1, the PCM model performed equally well than the RF models calculated for single transporter datasets. Interestingly, the individual model and the PCM model for GAT1 exhibit considerably lower performance (test and training set) than the models for the other three transporters. For the test set, this is predominantly due to one heavy outlier (compound **76**, prediction error = 1.9), which contains a photoswitchable azo moiety - a scaffold only present in the test set (see Figure 2c,d). For the training set, the poor cross-validation statistics could be caused by higher structural diversity within the GAT1 dataset compared to the other GATs as the 217 compounds comprise in total 85 Murcko scaffolds compared to 60 in mBGT1 (113 compounds), 75 in mGAT2 (163 compounds) and 78 in mGAT3 (179 compounds). The models for the other three transporters show good to excellent performance, especially for the test set (see Table 1).

**Table 1:** Overview of the performance of individual RF models and RF PCM models.

|  | 10-fold cross-validation |  | Test set |  | Bioactivities |  |
| --- | --- | --- | --- | --- | --- | --- |
|  | R2 | RMSE | R2 | RMSE | Training set | Test set |
| **mGAT1** | 0.53 | 0.78 | 0.63 | 0.52 | 217 | 22 |
| **mBGT1** | 0.67 | 0.43 | 0.81 | 0.26 | 113 | 22 |
| **mGAT2** | 0.68 | 0.43 | 0.91 | 0.38 | 163 | 22 |
| **mGAT3** | 0.73 | 0.39 | 0.85 | 0.49 | 179 | 22 |
| **all mGATs PCM** | 0.68 | 0.51 | 0.81 | 0.42 | 672 | 88 |
| **mGAT1 PCM** | 0.51 | 0.77 | 0.66 | 0.52 | 217 | 22 |
| **mBGT1 PCM** | 0.68 | 0.42 | 0.83 | 0.24 | 113 | 22 |
| **mGAT2 PCM** | 0.81 | 0.34 | 0.9 | 0.39 | 163 | 22 |
| **mGAT3 PCM** | 0.81 | 0.32 | 0.86 | 0.46 | 179 | 22 |

Analysis of the descriptor importance ranking showed that the top ranked ligand-based descriptors are general physicochemical parameters linked to polarizability (SMR), size (NumAtoms), and SlogP. With respect to the protein-based z3-scales descriptors, the amino acids in position 13 (L300/Q299/L294/L309 in mouse GAT1/BGT1/GAT2/GAT3, respectively) ranked highest (Figure 2b). Q299 is a unique residue for BGT1, corresponding to leucine in all other GATs (Figure 2). Strikingly, recent studies demonstrated that Q299 is a key residue relevant for activity and selectivity of BGT1 inhibitors such as 2-amino-1,4,5,6-tetrahydropyrimidine-5-carboxylic (ATPCA) analogs^25^ and also the biphasic compound SBV2-114^12^ by providing hydrogen bond interactions. Second and third ranked are amino acids which are unique in GAT1, while being conserved in the other subtypes such as the amino acids in position 1 (Y60/E52/E48/E66) and the amino acids in position 12 (G297/A296/A294/A311, see Figure 3). Mutational studies have already confirmed the relevance of E52 in hBGT1 (amino acid position 1, Figure 3) for the activity of ATPCA analogs^25^, underlying once more that the PCM models could identify critical features for activity and selectivity.

## Conclusion

PCM modeling has been shown to be a powerful tool for calculating QSAR models as it enables the utilization of structure and ligand-based information for predicting the activity profile of compounds towards a group of proteins. By analyzing the descriptor importance ranking, physicochemical properties of both ligands and amino acids can be derived that might be relevant for activity. In this case study, we generated for the first time a PCM model that predicts the activity of ligands for the four subtypes of the mGATs. Overall, the PCM model performs equally well compared to models calculated for each transporter individually. However, the PCM model allows to identify residues relevant for activity. Strikingly, the descriptor importance ranking of all protein descriptors identified the residues in the position L300/Q299/L294/L309 as well as Y60/E52/E48/E66 in mouse GAT1/BGT1/GAT2/GAT3 as most relevant. Both residue positions have been already confirmed by mutational studies to be important for the activity and selectivity of BGT1 and GAT3. Thus, we believe that proteochemometric modeling is a versatile tool with great potential for analyzing ligand-transporter interaction not only for the GATs but also for other SLC-transporter families.

## Acknowledgement

This work was supported by the Austrian Science Fund FWF (grant W1232).

## Notes

### Competing Interest Statement

The authors have declared no competing interest.

